# Theoretical analysis predicts an optimal therapeutic strategy in distinct parkinsonian landscapes of the striatum

**DOI:** 10.1101/2020.07.19.210690

**Authors:** Mathias L. Heltberg, Hussein N. Awada, Alessandra Lucchetti, Mogens H. Jensen, Jakob K. Dreyer, Rune N. Rasmussen

## Abstract

Parkinson’s disease (PD) results from a loss of dopaminergic neurons. The age of disease onset, its progression and symptoms vary significantly between patients, pointing to a complex relationship between neuron loss and PD etiology. Yet, our understanding of the clinical variability remains incomplete. Here, we use biophysical modelling to characterize the dopaminergic landscape in the healthy and denervated striatum. Based on currently proposed mechanisms causing PD, we model three distinct denervation patterns, and show notable differences in the dopaminergic network as denervation progresses. We find local and global differences in the activity of two types of striatal neurons depending on the denervation pattern. Finally, we identify an optimal cellular strategy for maintaining normal dopamine signaling when neurons degenerate stochastically within our model. Our results derive a conceptual framework in which the clinical variability of PD is rooted in distinct denervation patterns and forms testable predictions for future PD research.

## Introduction

Parkinson’s disease (PD) is a common neurodegenerative disorder, affecting 1% of people over the age of 60 worldwide^1^. The disease is caused by a progressive loss of dopaminergic neurons in the substantia nigra pars compacta (SNc)^2,3^, and symptoms typically emerge when 60–80% of these neurons are lost^4,5^. Notably, the age of onset, disease progression, response to treatment, and symptoms are highly variable between patients^6,7^, pointing to a complex relationship between neuron loss and PD etiology that remains to be understood.

Dopaminergic SNc neurons send projections to the dorsal striatum in the basal ganglia (Fig. 1a), an important area for motor function and executive control^8^. These projections promote movement by modulating the excitability of GABAergic striatal spiny projection neurons (SPNs) by activating D1- or D2-class dopamine (DA) receptors^9,10^. DA increases the excitability of D1 receptor-expressing SPNs (D1-SPNs) and decreases the excitability of D2 receptor-expressing SPNs (D2-SPNs)^8,11^. D1- and D2-SPNs are critical components of two distinct pathways, traditionally thought to control movements in opposing ways: the direct pathway promotes desired movements while the indirect pathway suppresses unwanted movements^8,12–14^ (Fig. 1b). In PD, dopaminergic neurons are progressively lost, leading to striatal DA depletion, abnormal SPN activity and movement deficits^3,13,15^. Despite the central role of failing DA signaling in PD etiology, little is known about the nature of striatal DA signaling before and during disease progression, posing a significant obstacle to the development of therapeutic strategies which maintain normal DA signaling in PD patients.

**Fig. 1.**
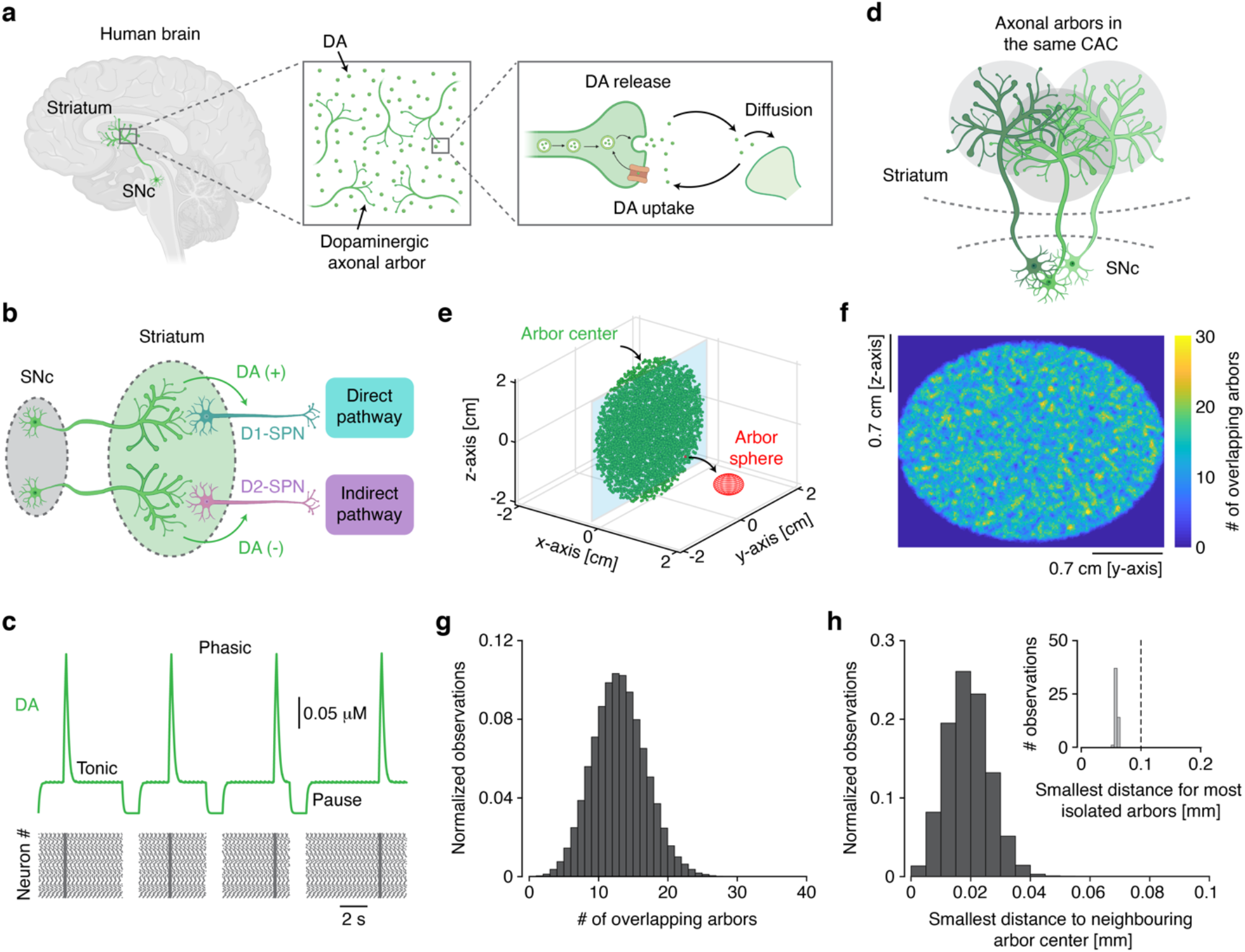
Functional and spatial characterization of DA signaling in the healthy human striatum. **a**, Diagram of dopaminergic innervation and signaling in the human striatum. **b**, Diagram of dopaminergic regulation of D1- and D2-SPNs, parts of the direct and indirect pathway, respectively. **c**, Trace showing DA signaling and the underlying dopaminergic neuronal firing pattern. **d**, Illustration of overlapping dopaminergic axonal arbors belonging to the same CAC. **e**, Visualization of dopaminergic axonal arbors in the striatum; each arbor center is marked with a circle. For visibility, only 10% of arbors are shown. Red sphere shows the area subsumed by an arbor from one neuron. Notice that all arbors belong to the same CAC, represented by them all having the same color. **f**, Heatmap of the distribution of overlapping arbors in the two-dimensional plane denoted in **e. g**, Distribution of the number of overlapping arbors for each individual arbor. **h**, Distribution of the smallest distance to the nearest neighboring arbor center for each arbor. Inset: smallest distance to nearest neighboring arbor center for the most isolated arbors found using Voronoi tessellation.

Efforts focused on understanding the molecular cascades underlying PD neurodegeneration^16^ have proposed different mechanisms, including the prion-hypothesis^17,18^ and oxidative stress^19,20^. However, little attention has been given to investigating the spatial and temporal patterns of dopaminergic neuron loss. Clinical imaging techniques, measuring DA transporter densities, provide a correlate of dopaminergic innervation^21,22^, but cannot resolve the fine-scale organization of neurons at cellular resolution. In animal models, neuronal firing and DA signals can be recorded invasively^23,24^ and correlated with dopaminergic neuron density postmortem. In addition to the challenge of being limited to a highly localized area, this approach lacks the temporal scale needed to track slow changes in neuron density and DA signaling.

Here, we developed biophysical models to probe cause-and-effect in a reduced parameter space. By characterizing the spatial patterns and processes of DA signaling degeneration, we found that distinct denervation patterns can be distinguished by unique temporal evolutions and DA signaling dynamics. Next, we show that these denervation patterns, through the effect of DA transients, differentially affect both the local and global activity of D1- and D2-SPNs. Finally, we use this to demonstrate that an ideal compensatory cellular strategy for maintaining normal DA signaling, despite neuron loss, is to enhance both DA release and DA uptake in parallel. These results support a conceptual framework where the clinical manifestations of PD are rooted in the distinct denervation patterns and, importantly, provide theoretical predictions to be experimentally tested.

## Results

### Functional and spatial characterization of DA signaling in the healthy striatum

We began our investigation by modelling DA signaling in the fully innervated human striatum, specifically the putamen, which we defined as the healthy state^25,26^. We simulated the firing of dopaminergic SNc neurons and described the DA concentration in the extracellular space. For this, we employed a model describing DA in a subvolume of 10^3^ μm^3^ (see Supplementary Information [SI]). Given the estimated density of ∼0.1 dopaminergic axonal terminals per μm^3^ in the healthy striatum^25–27^, this volume contains on average 100 terminals, each of which was treated as an individual element. This is a reasonable approximation since each time a neuron fires, only a small fraction of its terminals releases transmitter due to the probabilistic nature of this process. Estimates of the vesicular release probability of dopaminergic terminals are thus within the range of 6–20%^25,28^, and only ∼30% of the terminals may contain the molecular machinery for exocytosis^29^. Based on this, we employed a deterministic mean-field model that approximates DA inside the *i*’th subvolume as:

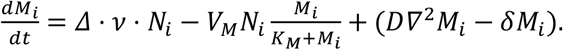

Here *M* is the DA concentration, *Δ* is the amount of DA released by a terminal, *ν* is the neuronal firing frequency, and *N* is the number of terminals within the subvolume. DA remains active in the extracellular space until it is removed by either transporters or degraded enzymatically (Fig. 1a), so we modeled transporter-mediated DA uptake after the Michaelis-Menten uptake equation. We also included a simple degradation term, and a term to account for the diffusion between neighboring subvolumes. However, for all but extreme cases, these are relatively negligible and therefore they are placed in parentheses. We assumed that each dopaminergic neuron can express one of two firing patterns: tonic or phasic firing^30,31^. From this, we obtained DA concentration time courses that clearly reflected the underlying neuronal firing patterns, exhibiting periods of tonic and phasic DA signaling, and pauses where DA is cleared from the extracellular space (Fig. 1c). Similar to naturally occurring DA transients measured in the striatum *in vivo*^32^, the maximal DA concentrations observed in our simulations were sub-micromolar (Fig. 1c).

We next characterized the dopaminergic innervation of the striatum at a macroscale. To mimic the shape of the putamen in the human striatum, we modeled it as an ellipsoid. Dopaminergic innervation was constructed by filling the volume with axonal arbors from 10^5^ SNc neurons, based on estimates from human SNc^33^ and the fact that these neurons have wide-spread projection targets^34^. Each neuron contributed with a spherical arbor with a radius of 0.5 mm, wherein the density of terminals was constant^27,35^. It should be noted that the experimental evidence for the exact volume of dopaminergic axonal arbors is still limited, especially in the human brain, but we consider the chosen value the best available estimate. To characterize the organization and spatial coverage of arbors within the striatum, we analyzed whether innervation was coherent within the striatum. This can be quantified by the number of what we termed *contiguous arbor classes* (CACs) — inspired by the mathematical analysis of communication classes. We assumed that dopaminergic neurons belonged to the same CAC if their arbors considerably overlapped (their arbor centers less than 0.5 mm apart; Fig. 1d). Hence, if the number of CACs is low, it suggests a high degree of spatial coverage and cohesion, and vice versa. For classifying neurons into CACs, we used a Markov chain-inspired algorithm (see SI). We found that all neurons in the healthy striatum belonged to the same CAC (Fig. 1e), suggesting a high degree of coverage (Fig. 1f). For each neuron, we also counted the number of overlapping arbors from other neurons, and this metric followed a Poisson distribution (Fig. 1g; see SI).

From the equation above, it is evident that decreasing *N*_*i*_ does not affect the steady state DA concentration considerably unless this value is approximately zero. From calculations on the diffusion equation (see SI), we determined that each point within the striatum with a distance larger than 0.1 mm to its nearest neighboring arbor was defined as isolated. We therefore searched for spatially isolated areas where the innervation was sparse, since such areas would be more susceptible to impairments in DA signaling during denervation. Using Monte Carlo simulations, we approximated the distribution of smallest distances and used Voronoi tessellation to find the most isolated points (Fig. 1h; see SI). This showed that no isolated areas existed in the fully innervated striatum of our model.

These results demonstrate that in the healthy striatum, DA exhibits three clearly distinct signaling periods: tonic, phasic and pauses. Furthermore, dopaminergic arbors comprise a network that densely covers the striatum, where no isolated areas exist.

### Different denervation patterns break down the dopaminergic network with distinct evolutions

In biology, structure often informs function. We therefore probed the spatial landscape of dopaminergic arbors in the denervating striatum. The cellular pathways involved in the loss of dopaminergic neurons are a fundamental question beyond the scope of this study. Instead, we sought to characterize the organization of the remaining innervation arising from distinct models of progressive neuron loss.

To describe denervation, we first assumed that all neurons have the same probability of dying, independent of their spatial position, and we termed this model *random denervation* (RD; Fig. 2a). The second model, *prion-like denervation* (PLD; Fig. 2b), was inspired by proposed mechanisms where protein aggregates spread between neurons and cause their degeneration^17,18,36^. A small set of neurons was initially infected, and at every timestep each infected neuron infected two neighboring neurons before being removed from the network. It should here be noted that whether this spread of toxic protein aggregates predominantly occurs between the somata in the SNc or between axons within the striatum is still unclear. Yet, given that dopaminergic neurons with nearby somata in the SNc seem to have nearby axonal arborizations in the striatum^34^, we consider our PLD model valid in both of these scenarios. In the third model, *stress-induced denervation* (SID), neurons die due to oxidative cellular stress^19,20^ (Fig. 2c). When dopaminergic neurons degenerate, the remaining neurons may upregulate their DA synthesis and firing activity in an attempt to maintain DA signaling. However, these neurons may already be close to their maximal metabolic capacity^37^, and increased activity could trigger stress-induced degeneration^19,20^. In this context, we assumed that the pace of neuronal death is a function of the number of remaining neighbors; neurons with few overlapping arbors have a higher risk of dying compared to neurons with many overlapping arbors. It is here important to establish, that we do not consider any of these three models as “ ground truth” mechanisms for the process of dopaminergic denervation, nor do we necessarily consider them mutually exclusive, but they each represent the simplest possible algorithm for studying the self-organization of these complex phenomena. For the purpose of our investigations, we therefore deem these distinct models valuable as a framework for testing the effects of different denervation patterns.

**Fig. 2.**
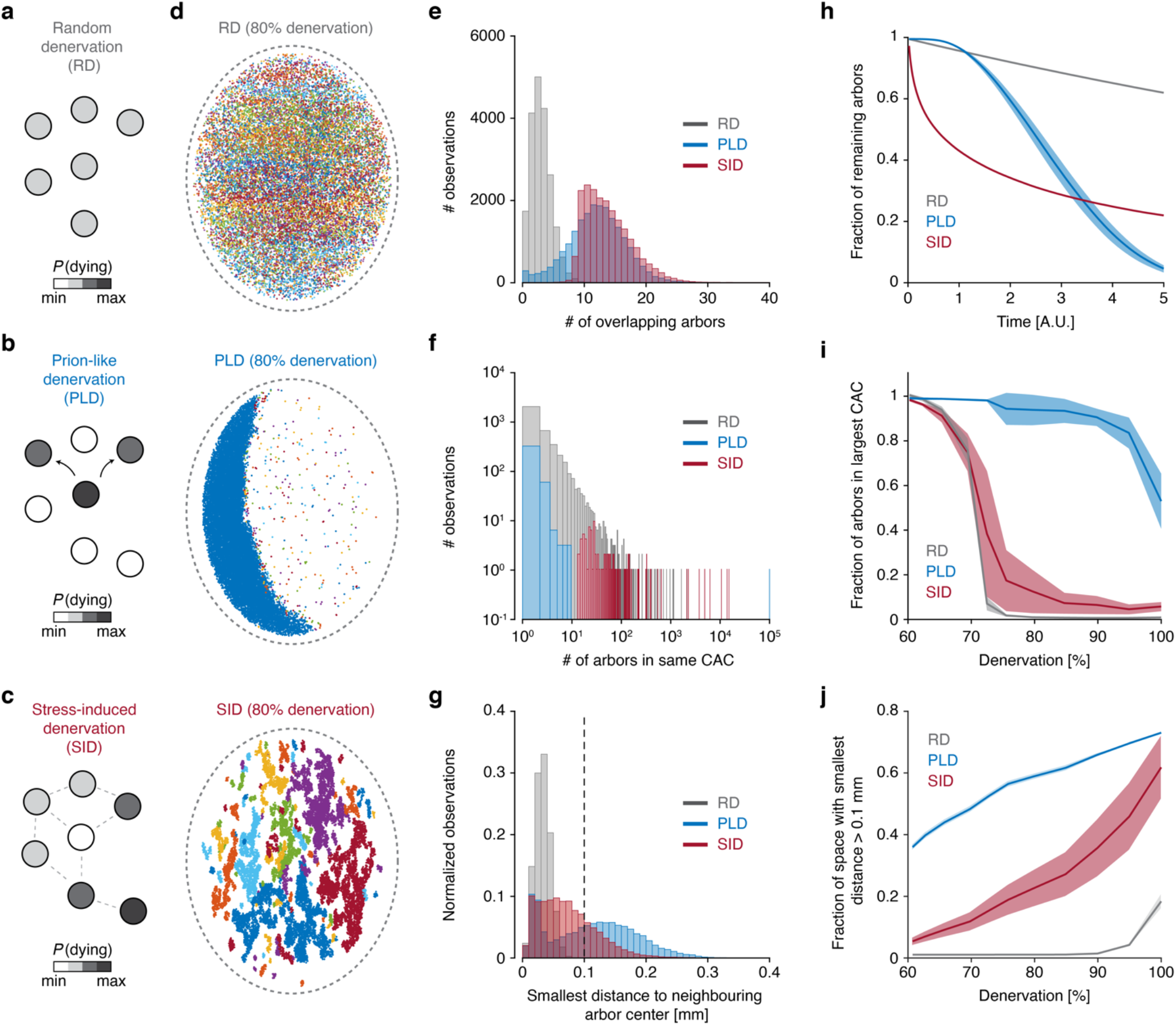
Different denervation patterns break down the dopaminergic network with distinct evolutions. **a**–**c** Diagrams of network mechanism for RD, PLD, and SID. The color of each dopaminergic neuron (circle) corresponds to probability of death. In **c**, dotted lines denote overlap of arbors. **d**, Visualization of the dopaminergic axonal arbor network following RD, PLD and SID. Colors correspond to separate CACs. **e**, Distributions of the number of overlapping arbors for each individual arbor. **f**, Distributions of the number of arbors in each CAC. **g**, Distributions of the smallest distance to the nearest neighboring arbor center for each arbor. Dotted line denotes threshold for classifying isolated areas. In **e–g**, denervation is 80%. **h**, Fraction of remaining arbors as a function of time. **i**, Fraction of arbors belonging to the largest CAC as a function of denervation. **j**, Fraction of striatal space with smallest distance to nearest arbor larger than 0.1 mm (isolated area) as a function of denervation. In **h**–**j**, full line is mean, and shading is standard deviation.

The three models resulted in distinct spatial landscapes, each characterized by a unique dopaminergic network breakdown (Figs. 2d–2g). For RD, the remaining arbors covered the entire striatal space but no longer belonged to the same CAC. In contrast, for PLD, large fractions of the striatum were deprived of arbors and instead dominated by one or two subregions with seemingly normal innervation. For SID, arbors were concentrated in small, isolated subregions, each forming its own CAC. We quantified these observations by the distribution of the number of overlapping arbors (Fig. 2e). For PLD, a notable fraction of arbors had very low numbers of overlapping arbors, whilst a larger fraction had numbers similar to those in the healthy striatum. In SID, only arbors with many overlapping neighbors remained. Importantly, a commonality of all models was that the dopaminergic network broke down into multiple CACs, but in distinct patterns (Fig. 2f): RD had only small classes remaining, PLD contained many small but also one dominating class, whereas SID contained many classes containing 100 or more arbors. We also assessed the emergence of isolated areas (Fig. 2g). For RD, no isolated areas existed. In contrast, for both PLD and SID, the striatum contained numerous isolated areas, deprived of arbors.

Next, we followed spatial characteristics as a function of denervation. First, we determined the percentage of remaining arbors as a function of time (Fig. 2h). For RD, this followed an exponential decay with a relatively slow temporal progression. Interestingly, for PLD, the curve followed a convex function, suggesting that neuron loss accelerated with time, whereas the curve for SID followed a concave function, indicating that denervation in this scheme started fast, but then slowed with time. These results have predictive strength and can be mathematically described by stretched exponentials of the form: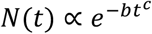, with *b* being the decay rate and *c* = 1 for RD, *c* > 1 for PLD and *c* < 1 for SID. We next characterized the breakdown of the spatial network, by calculating the fraction of arbors in the largest CAC (Fig. 2i). PLD kept one dominating class until the final stage of denervation, while RD and SID were characterized by a tipping point, at which the network dramatically transitioned from fully coherent to segregated into multiple classes. Interestingly, this transition occurred around 75% denervation, which often correlates with the onset of symptoms in PD patients^4,5,38,39^. Finally, we probed the emergence of isolated areas, practically devoid of DA signaling. At 75% denervation, isolated areas comprised ∼ 50% and 20% of the striatum in the PLD and SID models, respectively (Fig. 2j). At the same denervation level, no isolated areas existed for RD, but these emerged at severe denervation.

Overall, we found notable spatial and temporal differences between distinct models of dopaminergic denervation. These differences between models were remarkably robust to changes in key parameters, that is, axonal arbor volume and numbers of dopaminergic neurons in the healthy state (Supplementary Fig. 1).

### Dopaminergic denervation affects cAMP production and the activity of striatal SPNs

The excitability of SPNs is, along with several other factors, strongly regulated by DA^8–11^. Thus, we next asked how dopaminergic denervation affects the activity of individual SPNs. Previous work has shown that D1 and D2 receptors have low and high DA affinity, respectively^40^ (Fig. 3a). DA regulation of SPN excitability is mediated by the signaling molecule cyclic adenosine monophosphate (cAMP). D1 and D2 receptor activation increases and decreases the production of cAMP, respectively, and cAMP in turn regulates SPN ion channels^8– 10^. Inspired by previous work^26^, we described the intracellular cAMP concentration in D1- and D2-SPNs as:

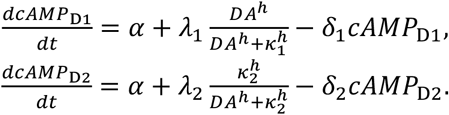

**Fig. 3.**
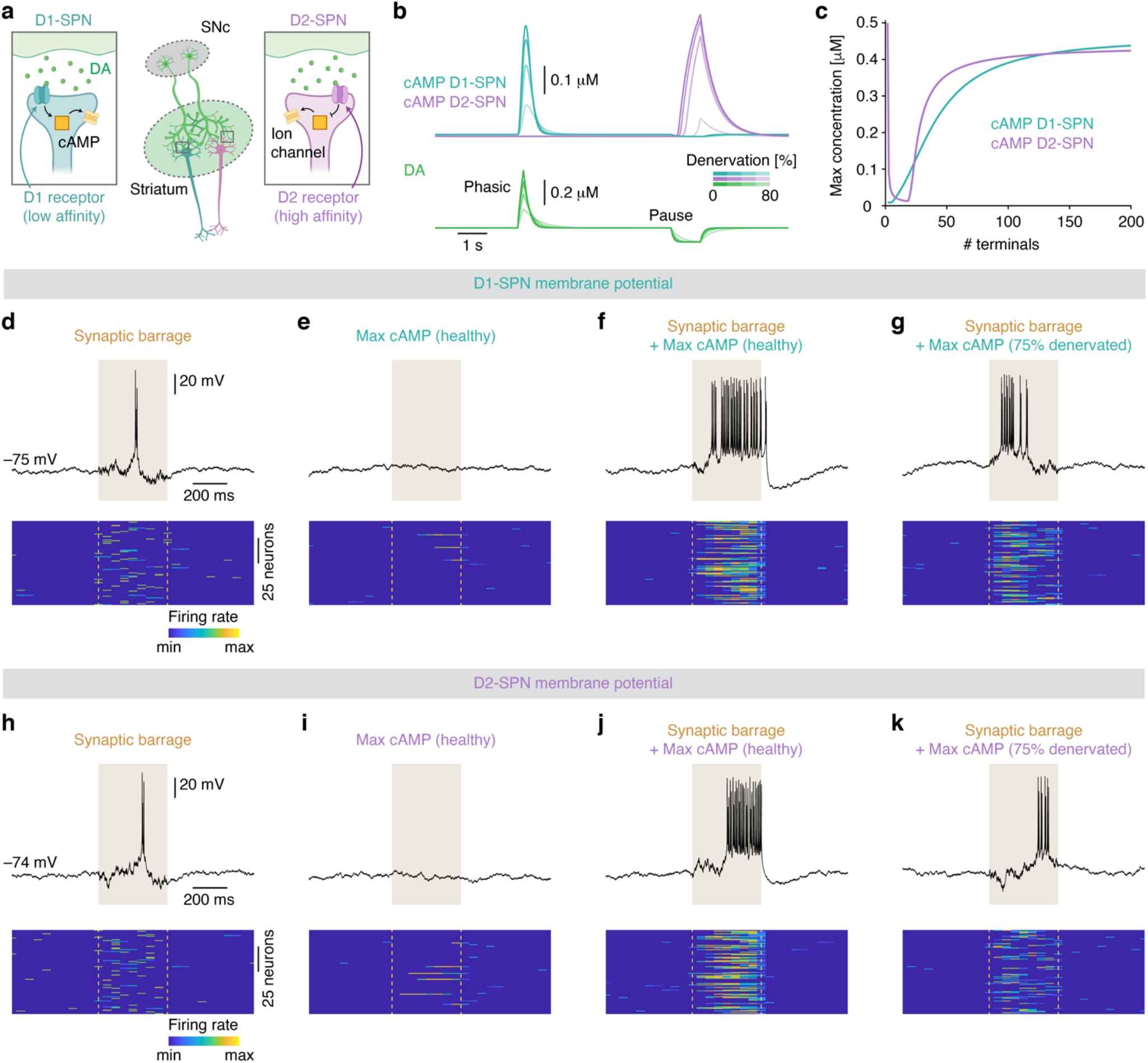
Dopaminergic denervation affects cAMP signaling and excitability of striatal SPNs. **a**, Diagram of how DA stimulates and inhibits the production of cAMP in D1- and D2-SPNs, respectively. **b**, Traces showing cAMP in D1- and D2-SPNs as a function of DA signaling. **c**, Maximal cAMP concentration in D1- and D2-SPNs during dopaminergic phasic firing and firing pauses, respectively, as a function of the number of dopaminergic terminals. **d**–**k**, Membrane potential dynamics of D1- and D2-SPNs in response to synaptic barrages (**d, h**), cAMP stimulation (**e, i**), synaptic barrages in combination with cAMP stimulation in the healthy state (**f, j**), or synaptic barrages in combination with cAMP stimulation in the 75% denervated state (**g, k**). Raster plots show the firing rate across time for 100 simulated neurons in each condition.

Here *α* is the steady state production of cAMP and *δ* is its spontaneous decay. In addition, receptor-dependent cAMP production was implemented: cAMP in D1- and D2-SPNs increased and decreased with DA stimulation, respectively. With increasing denervation, cAMP production during phasic firing became progressively lower in D1-SPNs, whilst in D2-SPNs it became progressively lower during firing pauses (Figs. 3b and 3c).

We next asked how these impairments in cAMP signaling may manifest in the activity of SPNs. For this, we used a Hodgkin-Huxley-inspired model — known as the Averaged-Neuron model^41,42^ — to simulate the membrane potential (V_m_) of D1- and D2-SPNs. This model contains extrinsic and intrinsic ion channel conductances. The extrinsic conductances are NMDA, AMPA, and GABA_A_ ion channels. The intrinsic conductances are voltage-gated and persistent Na_+_ channels, voltage-gated Ca_2+_ channels (Ca_V_), and voltage-gated, leak, fast A-type, inwardly rectifying, slowly inactivating (K_SI_), and Ca_2+_-dependent (K_Ca_) K_+_ channels_41,42_ (see SI). Using this model, we could closely mimic the V_m_ dynamics of SPNs. These neurons are characterized by their transitions between downstates, with a hyperpolarized V_m_, and upstates, with a plateau V_m_ depolarization upon which action potentials are fired^43,44^. In our model, upon increased levels of synaptic barrages (implemented by increasing the stochastic noise of the V_m_), both D1- and D2-SPNs transitioned into a brief upstate in which multiple action potentials were fired (Figs. 3d and 3h).

To implement the effect of DA, via its regulation of cAMP, on the V_m_ dynamics of D1- and D2-SPNs, we targeted the high-threshold Ca_V_ (N- and P-type), K_SI_, and K_Ca_ channels, which are negatively influenced by DA and cAMP signaling^10,11,45–48^. Thus, for increasing cAMP levels, the conductance of these channels decreases and vice versa (see SI). We note that other channels may also be subject to dopaminergic modulation, such as persistent Na^+^ or NMDA channels, but to limit the parameter space, we here focus on the abovementioned Ca^2+^ and K^+^ channels. For stimulating D1- and D2-SPNs, we used the cAMP concentrations observed during dopaminergic phasic firing and firing pauses, respectively. This was motivated by the result that, in the healthy striatum, the maximal cAMP production in D1- and D2-SPNs was observed during these two phases respectively (Fig. 3b). In itself, cAMP stimulation was very rarely sufficient to evoke a transition from the downstate to an upstate in types of SPNs (Figs. 3e and 3i), supporting the notion of DA as a “ modulator” rather than a “ driver” ^9–11^. However, if DA and cAMP stimulation coincided with increased levels of synaptic barrages, this triggered a robust upstate that lasted longer and elicited more action potentials than with synaptic barrages alone (Figs. 3f and 3j); this demonstrates that cAMP powerfully regulates the excitability of D1- and D2-SPNs. As a result, in the denervated state, the activity of SPNs was notably affected (Figs. 3g and 3k): the duration of the upstate and the firing rate during the upstate were strongly diminished in both D1- and D2-SPNs. These findings were largely replicated using the far simpler Izhikevich model (Supplementary Fig. 2a; see SI), suggesting model-invariance.

Together, these data demonstrate that DA signaling, via its downstream effector cAMP, potently regulates the firing activity of SPNs, and this regulation is impaired in the denervated striatum.

### Distinct denervation patterns differentially affect global SPN firing activity

Next, we sought to investigate how different denervation patterns affect the activity of D1- and D2-SPNs across striatal space. For this, we spatially mapped the maximal D1-SPN firing rate during dopaminergic phasic firing, and maximal D2-SPN firing rate during firing pauses, determined in the Averaged-Neuron model (Figs. 4a and 4b). In RD, although almost all the subregions had relatively low DA levels compared with the healthy striatum, this was still sufficient to evoke intermediate D1-SPN firing rates across the extent of the striatum. In contrast, in both PLD and SID, D1-SPN firing was high only in the subregions with preserved DA innervation. Noticeably, PLD transformed the striatum into a strongly polarized activity map, whereas SID caused local heterogeneity. For D2-SPNs, the emergence of isolated areas, resulted in a very different outcome. Since the maximal DA concentration in isolated areas is zero (except for small diffusive fluctuations), D2-SPN firing rates were here very high, most profoundly expressed for PLD and SID. We note here that, under physiological conditions, D2-SPNs residing in regions deprived of DA signaling might adapt by downregulating their firing rates to maintain homeostasis. In this scenario, the results would likely be similar to those for D1-SPNs.

**Fig. 4.**
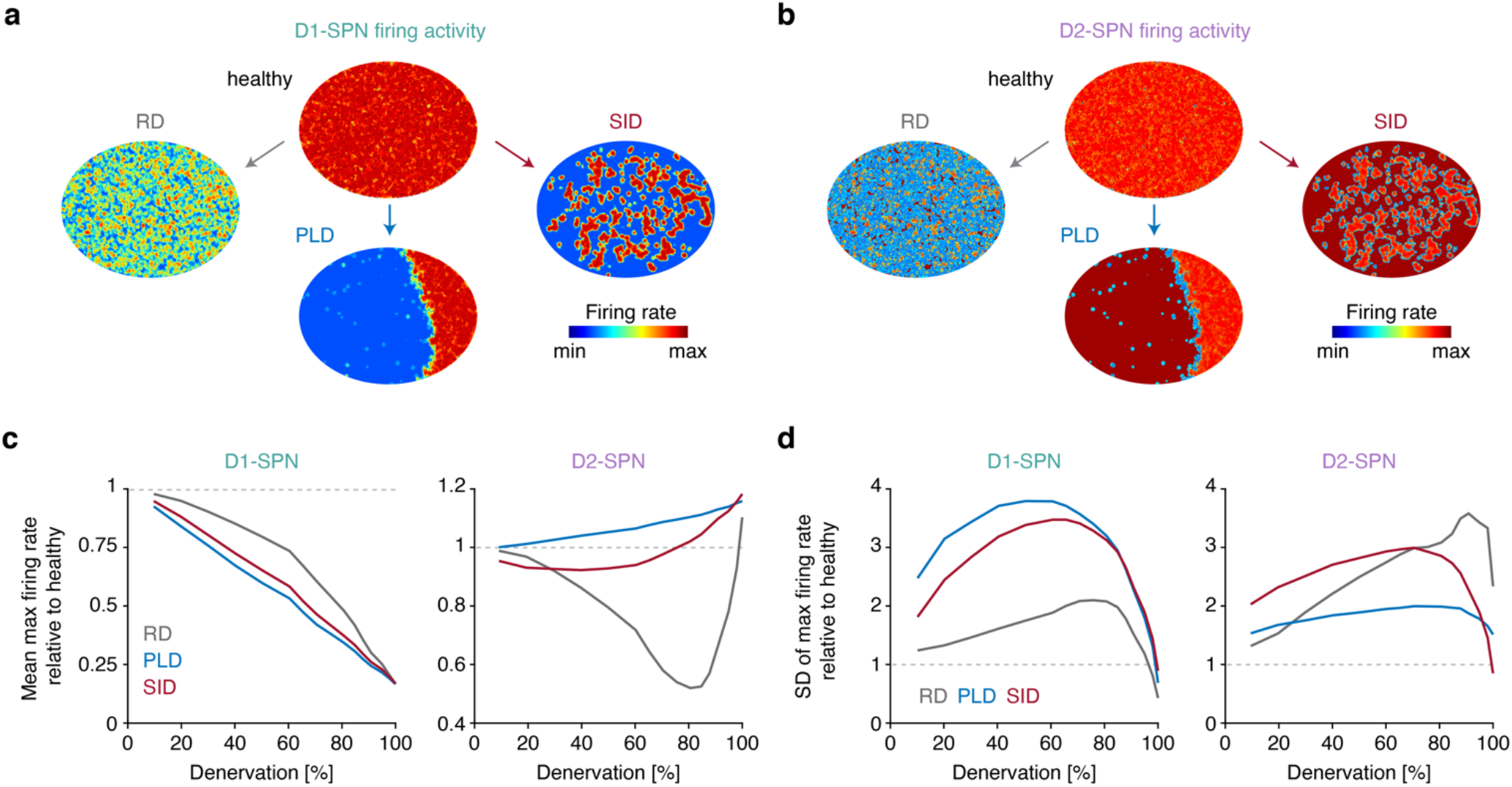
Distinct denervation patterns differentially affect global striatal SPN firing activity. **a**–**b** Maximal firing activity of D1- and D2-SPNs across space in the healthy and 75% denervated striatum for the three denervation patterns. **c**–**d** Spatial mean and standard deviation of maximal firing activity in D1- and D2-SPNs as a function of denervation.

Finally, we characterized SPN activity in the three denervation models as a function of denervation. The mean D1-SPN firing rates decreased linearly as a function of denervation in all models (Fig. 4c). We also noted that the standard deviation of D1-SPN firing was smaller in RD compared to both PLD and SID, indicating spatial homogeneity of firing levels (Fig. 4d). The effect on mean D2-SPN firing rates was different: firing increased notably for PLD and slightly for SID, as a function of denervation (Fig. 4c). In RD, the firing rates decreased until it reached a minimum around 80% denervation, whereafter it rapidly increased. This observation is explained by the occurrence of isolated areas, deprived of DA signaling, resulting in a dramatic increase in cAMP production in D2-SPNs (Fig. 3c), in turn resulting in a profound increase in excitability. Comparing the three models, it is clear that the early progression of denervation (up to ∼60%) resulted in increased standard deviation of SPN firing rates for all denervation patterns (Fig. 4d). This increase in activity variance across neurons may thus be a fingerprint of the denervating striatum. All of the described results were fully replicated with the Izhikevich model (Supplementary Fig. 2).

Overall, these results show that the global firing activities of D1- and D2-SPNs are strongly affected by the specific spatial pattern of dopaminergic denervation.

### A dual presynaptic compensation strategy preserves DA signaling in the denervated striatum

Given that dopaminergic neurons loss may trigger compensatory mechanisms in the remaining neurons in an attempt to maintain normal DA signaling^49,50^, we sought to probe the potency of such mechanisms, in order to attempt predicting ideal therapeutic strategies. We included three presynaptic compensatory mechanisms in our model and tested their impact on DA signaling. First, remaining dopaminergic terminals may increase their DA release capacity^50–52^. We refer to this as *enhanced release compensation* (ERC; Fig. 5a):

**Fig. 5.**
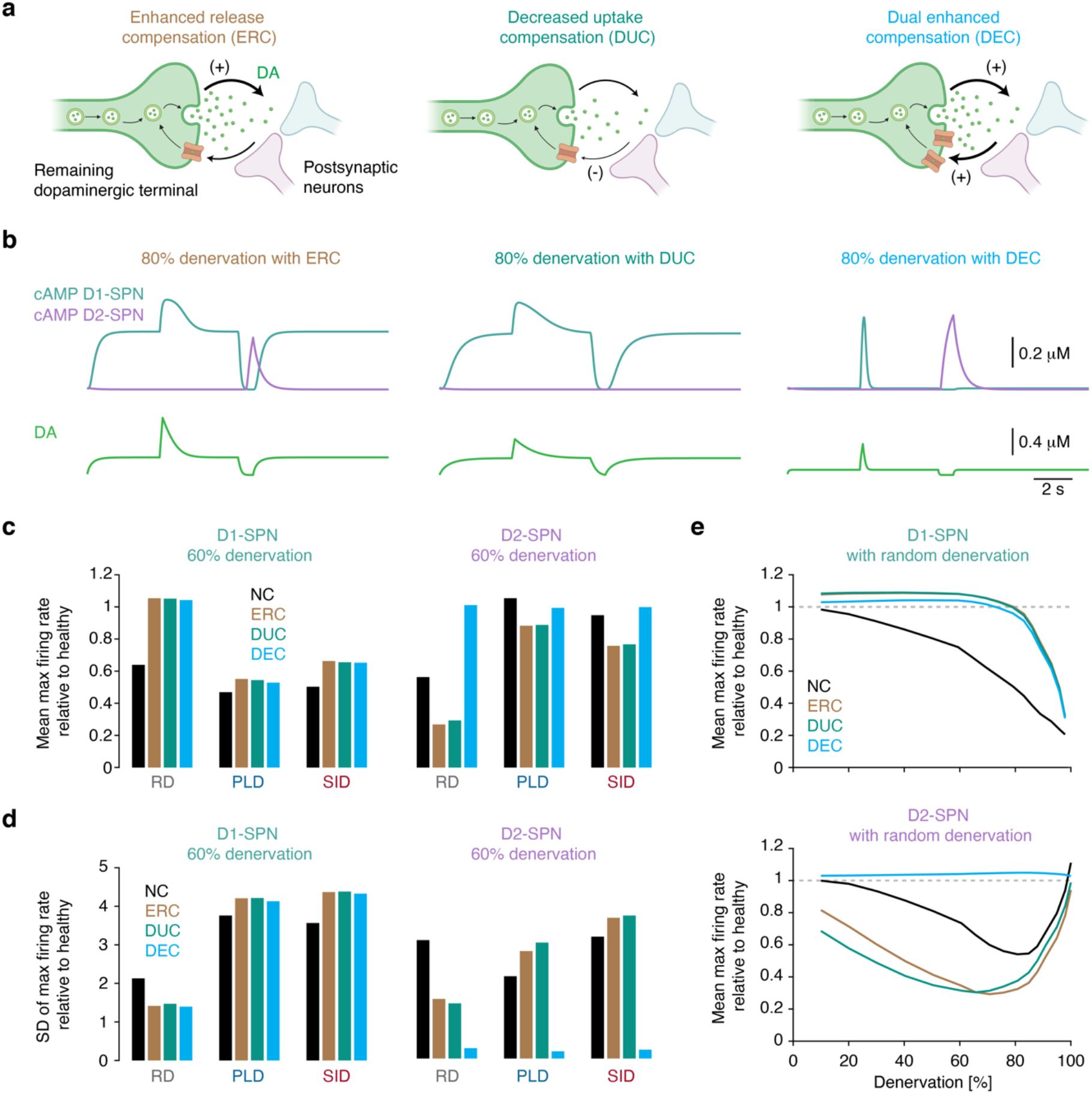
A dual presynaptic compensation strategy preserves DA signaling in the denervated striatum. **a**, Diagrams of mechanisms of the ERC, DUC, and DEC models. **b**, Traces showing cAMP in D1- and D2-SPNs as a function of DA signaling at 80% denervation in the compensation models. **c**–**d** Spatial mean and standard deviation of maximal firing activity in D1- and D2-SPNs as a function of denervation pattern and compensation model. **e**, Spatial mean of maximal firing activity in D1- and D2-SPNs as a function of denervation and compensation model in the randomly denervated striatum.

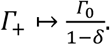

Here we have introduced the compensation parameter δ, a sigmoidal function going from zero to one as a function of the number of dopaminergic arbors covering a small volume. The parameter *Γ*_0_ refers to the DA release in healthy subregions, whereas *Γ*_+_ is the compensated release value. Second, DA transporters, expressed on terminals, may reduce their uptake capacity^5,50–52^. We refer to this as *decreased uptake compensation* (DUC; Fig. 5a):

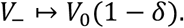

As above, the parameter *V*_0_, refers to the uptake value in healthy subregions, whereas *V*_–_ is the compensated uptake strength. Finally, we suggest a mechanism where neurons compensate by enhancing both DA release and uptake capacity in the terminals. Such a compensatory mechanism has not previously been suggested, and we refer to this as *dual enhanced compensation* (DEC; Fig. 5a); this is included in the model through changes in both the uptake and release parameters:

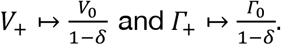

Here all parameters are defined as above.

With these implementations, we simulated DA signaling and the corresponding cAMP production in D1- and D2-SPNs with 80% denervation. As shown above, in the absence of any compensatory mechanism, DA release during tonic firing is unaffected, but notably affected during phasic firing and firing pauses (Figs. 3b and 3d). Here, during tonic firing, the DA concentration notably increased for ERC and DUC models; during phasic firing, DA was increased for ERC, and during firing pauses, DA removal was incomplete for DUC (Fig. 5b). Importantly, the DEC model preserved DA levels during both tonic and phasic firing at comparable levels to in the healthy state, while still allowing complete DA removal during firing pauses (Fig. 5b). We also tested several postsynaptic compensatory mechanisms previously proposed in the literature, including increased D2 receptor expression^53^, enhanced D1 and D2 receptor sensitivity^54^, and suppressed cAMP degradation in D1 and D2-SPNs^55^. These mechanisms were inadequate to restore DA and cAMP signals in SPNs (Supplementary Fig. 3), and we therefore did not explore these further.

Next, we asked if any of the presynaptic compensation mechanisms were able to ameliorate the impairments in SPN firing activity in the denervated state (Figs. 3 and 4). For this we calculated the spatial mean and standard deviation of the maximal D1- and D2-SPN firing rates in the three denervation models and combined these with the presynaptic compensation mechanisms (Fig. 5c). Interestingly, in the scenario of RD, only the DEC model preserved the mean level of D1- and D2-SPN firing rates. In contrast, in the case of PLD and SID, neither of the three compensation models were able to counteract the drastic decrease in D1-SPN firing with denervation, whilst all models performed relatively well for D2-SPN firing activity. For the standard deviation of the D1-SPN firing rates, we note that in the RD scenario, all compensation models, as well as the no compensation model, maintained this measure near the healthy level (Fig. 5d). In contrast, for PLD and SID, the standard deviation was notably increased for all compensation models, and curiously, the non-compensated model was most similar to the healthy state. For the standard deviation of the D2-SPN firing activity, none of the compensation models truly maintained this measure close to the healthy level, regardless of the denervation pattern (Fig. 5d). It is here worth noting that the DEC model, across all denervation patterns, maintained the standard deviation of D2-SPN firing activity at a very low level. This is because, in regions with low dopaminergic coverage, DA signaling from remaining terminals in the DEC model can compensate optimally, restoring coherent neuronal activity. The low standard deviation in firing activity across neurons means that all striatal subregions are capable of generating a very similar firing response upon dopaminergic stimulation. Overall, from these results, we conclude that the DEC model, in combination with the RD pattern, best preserved the global SPN firing activity, despite considerable denervation. In the final set of simulations, we thus explored this for all levels of denervation (Fig. 5e). For the DEC model, the firing rates of both D1- and D2-SPNs remained remarkably close to the healthy state, despite reaching severe denervation.

In contrast, for the ERC and DUC models, even at relatively low denervation, SPN firing differed from the healthy state. Specifically, around 60% denervation, the firing rates of D1- and D2-SPNs was notably higher and lower, respectively, compared to the healthy state. Therefore, the ERC and DUC mechanisms do not seem ideal as therapeutic strategies. These findings were generally replicated with the Izhikevich model (Supplementary Fig. 4), bolstering the robustness of these results.

Taken together, these results show that an ideal strategy to maintain normal SPN activity is to locally introduce a dual compensation mechanism — increasing both DA release and uptake capacity — and to globally minimize the dopaminergic arbor density differences, or at least avoid the emergence of isolated areas.

## Discussion

In this work, we used biophysical modeling to investigate the spatial and functional landscape of dopaminergic signaling in the healthy and parkinsonian striatum. First, we showed that the spatial pattern of dopaminergic denervation profoundly affects the structural and temporal breakdown of the dopaminergic network in the striatum. Second, we derived how the local and global activity of D1- and D2-SPNs were differentially affected as a function of the spatial dopaminergic denervation pattern. Third, we identified that the optimal cellular strategy for maintaining normal striatal DA signaling, when neurons are progressively lost, is to enhance both DA release and uptake capacity.

### Clinical variability in PD patients may be mediated by different dopaminergic denervation patterns

PD symptoms often present when 60–80% of dopaminergic neurons are lost^4,38^. Still, the age of onset, disease progression and symptoms can vary notably between patients^6,7^. We argue that the spatial pattern of denervation, caused by different molecular mechanisms, could be a central determinant for much of this clinical variability. Importantly, the shape of the denervation curves varied notably depending on the denervation mechanism (Fig. 2). After an initial slow denervation rate, the loss of neurons accelerated with time in the PLD model. In contrast, in the SID model, denervation slowed with time after an initial rapid loss of neurons. The temporal evolution of the PLD and SID models correlates remarkably well with the clinical progression pattern seen in the early and middle stages of PD^56,57^. Our model thus predicts that the dopaminergic denervation pattern may be a central determinant for the disease progression variability and etiology seen in PD. In patients with RD, disease progression may be slow, whereas in patients with PLD, the progression may accelerate rapidly. Hence, if the total density of striatal dopaminergic terminals can be measured as a function of time in the early stages of the disease, it may be possible to determine the molecular mechanism and denervation pattern causing PD in individual patients. Furthermore, it may be feasible to, at least in part, predict the disease progression time course, and from that determine the ideal therapeutic strategy for the individual patient. We therefore propose that future clinical experiments aim to measure the density of dopaminergic terminals in the striatum of PD patients over time, using for example single-photon emission computed tomography^21,22^, and to correlate this to disease progression. Combining the results from such experiments with biophysical modelling would elucidate the molecular and network mechanisms causing PD and disease variability. We also found that the absolute time course of dopaminergic denervation was remarkably distinct between the different denervation patterns (Fig. 2), and this observation could potentially aid clinicians in determining the differential diagnosis of parkinsonism. Clinical imaging of early-stage PD patients has shown that structural innervation differences in the striatum, albeit embracing a notably larger striatal area than our results, relates to different PD-related diseases. For example, large-scale asymmetry in striatal dopaminergic innervation associates with idiopathic parkinsonism^58,59^, while large-scale symmetric denervation associates with atypical parkinsonian syndromes such as supranuclear palsy^58,60–62^. This difference might be explained by the three denervation patterns described in our work. Thus, the denervation curves from our model, in combination with high-resolution imaging, might portent a valuable tool for distinguishing between different forms of parkinsonism in individual patients.

### Distinct dopaminergic denervation patterns may differentially affect the direct and indirect pathway

When dopaminergic neurons were lost, the burst firing during upstates of D1- and D2-SPNs was severely impaired (Fig. 3). Given that D1- and D2-SPNs are critical components of the direct and the indirect pathways, respectively, it is plausible that these two pathways would be affected as a result. The reduction of firing in D1-SPNs during phasic dopaminergic firing may complicate the initiation of voluntary movements, whilst the impaired firing in D2-SPNs during dopaminergic firing pauses may facilitate unwanted, involuntary movements. Given that, in the denervated striatum, DA signaling and SPN activity varied across space depending on the denervation pattern, we expect that different subregions of the striatum will have normal and abnormal activity of the direct and indirect pathways, depending on the denervation pattern. This may, at least in part, explain why disease symptoms can vary notably between PD patients. We mention this with the caveat that our simulations of SPN activity have their limitations. For example, DA is not the only modulator of SPN activity; also local GABAergic and cholinergic interneurons regulate the activity of SPNs^9,63^, and our simulations do not account for that. Furthermore, we here investigated the acute effects of dopaminergic denervation and our results do thus not take into account the long-term changes in glutamatergic synaptic activity that may develop in the striatum as a result of denervation^8,10^, nor did we explore possible changes in dopaminergic autoregulation^64^. Future work should aim to investigate the interplay between different dopaminergic denervation patterns and these other mechanisms regulating SPN activity, for example in a more comprehensive basal ganglia network model, as recently developed^65^.

### A dual cellular strategy maintains normal DA signaling and may delay severe symptoms in PD patients

The most common pharmacological treatment for PD is to administer levodopa, with the goal of increasing DA levels within the brain^66–68^. However, not all patients respond well to this treatment, and some experience side adverse effects with long-term treatment^66–68^. During the early stages of PD, the dopaminergic neuron loss is believed to be counterbalanced by endogenous compensatory mechanisms^49,50^. Knowledge of such mechanisms could reveal potential targets for novel therapeutic strategies^49^. We found that DA signaling cannot be fully characterized by only the tonic level. The peaks during phasic firing and the complete removal of DA during firing pauses are also an integral part of what we consider normal DA signaling, and likely play important roles in proper neuronal signaling. Therefore, when evaluating the therapeutic potential of a cellular target, it is important to assess its effects on the full DA signaling spectrum. Interestingly, the compensation strategy that best maintained DA signaling, despite severe denervation, was a dual mechanism that enhanced both the release and uptake of DA in the remaining neurons. In contrast to other mechanisms, this dual mechanism preserved the DA signaling spectrum, without increasing tonic DA levels. Thus, as long as some neurons remain, this mechanism is able to restore the DA signaling properties, emphasizing the great importance of avoiding areas completely devoid of DA terminals. This suggests that therapeutic strategies that achieve simultaneous enhancement of DA release and uptake capacity should be pursued. We hypothesize that this dual strategy might postpone the onset of severe symptoms by upholding normal DA signaling, and may also cause less side effects since baseline DA is maintained at a comparable level to that in the healthy striatum.

Our work constitutes a new conceptual model for the clinical manifestations of PD, while providing a clear set of theoretical predictions and testable hypotheses. We regard our biophysical modeling as the first step towards further experimental investigations required to test our results. We expect that these findings will lay the groundwork for new research directions within both basic and clinical sciences, aimed at better understanding and treating PD.

## Materials and Methods

This work uses numerical and analytical mathematical methods to theoretically investigate and characterize the dopaminergic innervation of the human striatum, and the networks that break down following denervation. In the first part of the paper, we use mean-field theory to derive a differential equation describing DA signaling in a mesoscopic region of the striatum. By inducing three distinct periods of dopaminergic neuron firing, we solve this numerically. Following this, we introduce individual axonal arbors in the striatum, using the random generator applied in MATLAB, calculate CACs, and estimates the unoccupied space by placing 10,000 random points in the striatum. Next, we repeat these measures for the three denervation mechanisms (RD, PLD, and SID). All denervation models are generated based on the described algorithms and simulated in MATLAB. We then introduce a differential equation describing cAMP following DA stimulation, which we solve numerically using MATLAB. With this, we record the maximal cAMP value for both D1- and D2-SPNs. We next introduce the Averaged-Neuron model and simulated this in Python using the Numba package. Using this, we measured the number of elicited action potentials in a short temporal window, in which the D1- and D2-SPNs were stimulated with the corresponding maximal cAMP level. This was done for 100 neurons at several different levels of denervation. Finally, we used the average number of action potentials recorded in this window as input for 10,000 randomly positioned points in the striatum, dependent on the denervation model and levels. Using these numbers, we calculate a spatial average and standard deviation of the maximal firing rates of D1- and D2-SPNs. A detailed description of all mathematical derivations and formulations, biophysical models, and algorithms are included in the SI appendix. Figures were assembled using Illustrator (Adobe), and schematic diagrams created using BioRender.

## Acknowledgements

We thank Ubadah Sabbagh, Akihiro Matsumoto and Eric Nicholas for critical comments on the manuscript. M.L.H. is grateful to Aleksandra Walczak and Thierry Mora for scientific discussions and valuable support. M.L.H. thanks Angela Taddei and Judith Mine-Hattab for encouragement and a fantastic scientific environment. M.L.H. and H.N.A. would like to thank Lene Oddershede for inspiration in the early stages of the project. R.N.R. acknowledges support from the Lundbeck Foundation (R230-2016-2326); M.L.H. and M.H.J. acknowledge support from the Danish Council for Independent Research and StemPhys DNRF Center of Excellence (DNRF116).

## Author contributions

M.L.H., H.N.A., M.H.J, J.K.D., and R.R. designed the research; M.L.H., and A.L. performed the research; M.L.H., H.N.A., A.L., J.K.D., and R.R. analyzed and interpreted the data; M.L.H., and R.R. made the figures; M.L.H., H.N.A., M.H.J., J.K.D., and R.R. wrote the paper.

## Data and code availability

Code is made publicly available at the GitHub repository: https://github.com/Mathiasheltberg/Theoretical_Denervation_ParkinsonsModel.

