## Supplementary Information for "Theoretical analysis predicts an optimal therapeutic strategy in distinct parkinsonian landscapes of the striatum"

##### Supplementary inventory

1. Models and methods
   - Simulations in the mean field model
   - Analytical derivation of the dopamine concentration
   - Simulation of the spatial dopaminergic network
   - Distribution of overlapping axonal arbors
   - Concentration of dopamine inside an isolated area
   - Voronoi tessellation
   - Defining contiguous arbor classes
   - Removing dopaminergic neurons
   - Averaged-Neuron SPN model
   - Izhikevich SPN model
   - Postsynaptic compensatory mechanisms
2. Figures
   - Supplementary Fig. 1
   - Supplementary Fig. 2
   - Supplementary Fig. 3
   - Supplementary Fig. 4
3. Tables
   - Supplementary Table 1

### 1 Models and methods

#### Simulations in the mean field model

To construct the mean field model, we considered a cubic volume of dimensions $(10 \times 10 \times 10)\,\mu\mathrm{m}^3 = 1000\,\mu\mathrm{m}^3$. Since the density of dopaminergic axonal terminals is estimated to be $0.1\,\mu\mathrm{m}^{-3}$, this volume contained 100 terminals. Next we considered the diffusion coefficient being $D \approx 380\,\mu\mathrm{m}^2\,\mathrm{s}^{-1}$ and thus the time for DA to equilibrate would be $\Delta t \approx \frac{\Delta x^2}{2D} \approx 0.03\,\mathrm{s}$ with $\Delta x = 5\,\mu\mathrm{m}$. We therefore performed the simulations with this timestep and assumed that a release of DA would affect the DA level in the entire volume. We also assumed that the influx from neighboring axonal arbors would be equal to the outflux, since the density of dopaminergic terminals would be homogeneous on these small length scales.

#### Analytical derivation of the dopamine concentration

As explained in the main text, the dopamine dynamics in the i-th subvolume of the striatum can be approximated as

$$\dot{M_i} = \Delta \cdot N_i - V_M N_i \frac{M_i}{M_i + K_M}, \tag{1}$$

if we neglect the natural degradation and the diffusion between neighboring subvolumes. The above differential equation is solvable, so we wished to describe $M_i(t)$. Dropping the i subscript for simplicity of notation, and defining $\Delta \cdot N - V_M N \equiv \gamma$ and $\Delta \cdot N K_M \equiv \delta$ we can write

$$\frac{dM}{dt} = \frac{M\gamma + \delta}{M + K_M} \Rightarrow \int \frac{M + K_M}{M\gamma + \delta} dM = t + C. \tag{2}$$

The integral can be solved, such that

$$\begin{aligned}
\int \frac{M + K_M}{M\gamma + \delta} dM &= \int \frac{M}{M\gamma + \delta} dM + \int \frac{K_M}{M\gamma + \delta} dM \\
&= \frac{K_M}{\gamma} \int \frac{1}{u} du + \frac{1}{\gamma^2} \int \frac{u - \delta}{u} du \qquad \text{with } u = M\gamma + \delta \\
&= \frac{K_M}{\gamma} \ln(u) + \frac{1}{\gamma^2} u - \frac{\delta}{\gamma^2} \ln(u) + C \\
&= \frac{M}{\gamma} + \ln(M\gamma + \delta) \cdot \left( \frac{K_M}{\gamma} - \frac{\delta}{\gamma^2} \right).
\end{aligned} \tag{3}$$

From this we can rewrite the equation

$$(\gamma M + \delta) + \ln(M\gamma + \delta) \cdot (-K_M V_M) = \gamma^2 t + \delta - C. \tag{4}$$

By setting $T \equiv \gamma M + \delta$, $-K_M V_M \equiv \tilde{\Delta}$ and $\gamma^2 t + \delta \equiv f(t)$ we obtain

$$\begin{aligned}
&T + \tilde{\Delta}\ln(T) = f(t) - C \\
&T + \tilde{\Delta}\ln\left(\frac{T}{\tilde{\Delta}}\right) = f(t) + \tilde{\Delta}\ln\left(\frac{1}{\tilde{\Delta}}\right) - C \\
&\frac{T}{\tilde{\Delta}} + \ln\left(\frac{T}{\tilde{\Delta}}\right) = \frac{f(t)}{\tilde{\Delta}} + \ln\left(\frac{1}{\tilde{\Delta}}\right) - C \\
&e^{T/\tilde{\Delta}}\frac{T}{\tilde{\Delta}} = \frac{e^{\frac{1}{\tilde{\Delta}}(f(t)-C)}}{\tilde{\Delta}} \\
&T = \tilde{\Delta} W\left(\frac{e^{\frac{1}{\tilde{\Delta}}(f(t)-C)}}{\tilde{\Delta}}\right)
\end{aligned} \tag{5}$$

with W being the Lambert W function. By inserting the initial conditions and rewriting the expression in terms of $M(t)$ we end up with

$$M(t) = \frac{1}{\gamma}\left[W\Big(e^{\frac{f(t)}{\tilde{\Delta}}} e^{\frac{M_0\gamma}{\tilde{\Delta}}} \frac{(\gamma M_0 \delta)}{\tilde{\Delta}}\Big)\tilde{\Delta} - \delta\right]. \tag{6}$$

#### Simulation of the spatial dopaminergic network

To construct the dopaminergic network, we considered each dopaminergic axonal arbor to be a sphere with radius $r_d = 0.5\,\mathrm{mm}$, centered at $(c_x, c_y, c_z)$, which are random coordinates that satisfy

$$\left(\frac{c_x}{a}\right)^2 + \left(\frac{c_y}{b}\right)^2 + \left(\frac{c_z}{c}\right)^2 \leq 1.$$

Here $a = 0.3\,\mathrm{cm}$, $b = 1.5\,\mathrm{cm}$ and $c = 2.1\,\mathrm{cm}$ correspond to the principal axes of the ellipsoid that model the striatum, whose volume is therefore $V_T = 4\pi/3\,(0.3 \times 1.5 \times 2.1)\,\mathrm{cm}^3 = 3.96\,\mathrm{cm}^3$. Having placed these spheres, we generated a connected network, where we let two dopaminergic neurons interact if the distance between the two arbor centers was less than or equal to the radius of one arbor.

#### Distribution of overlapping axonal arbors

We described the distribution of dopaminergic axonal arbors covering an arbitrary point in the striatum. For the healthy state, we estimated this analytically. We considered a region $\epsilon$ with radius $r_\epsilon << r_d$, centered at position $c_\epsilon$. If we randomly embedded an axonal arbor, with volume $V_d = \frac{4}{3}\pi r_d^3$, the probability that $\epsilon$ would be covered by this arbor would be

$$P(\epsilon \in V_d) = \frac{V_d}{V_T} \approx 1.25 \times 10^{-4}. \tag{7}$$

Assuming that all axonal arbors are embedded independently, we calculated the probability that a region in space $\epsilon$ is covered by n arbors, given that there are $N_T$ in total:

$$P(\{\Delta_{i\to\epsilon} \leq r_d\} = n | N_T) = \left(\frac{V_d}{V_T}\right)^n \left(1 - \frac{V_d}{V_T}\right)^{N_T - n} \frac{N_T!}{n!(N_T - n)!}. \tag{8}$$

Here $\{\Delta_{i\to\epsilon} \le r_d\}$ is the set of arbors with their center closer to $\epsilon$ than $r_d$. Rewriting $\frac{V_d}{V_T} \Rightarrow \frac{\frac{V_d}{V_T} N_T}{N_T}$, and given that $N_T >> n$, we made the following approximations,

$$\left(1 - \frac{\frac{V_d}{V_T} N_T}{N_T}\right)^{N_T - n} \approx \left(1 - \frac{\frac{V_d}{V_T} N_T}{N_T}\right)^{N_T} = e^{-\frac{V_d}{V_T} N_T} \text{ and } \quad \frac{N_T!}{(N_T - n)!} \approx {N_T}^n. \tag{9}$$

Inserting these we end up at

$$P(\{\Delta_{i\to\epsilon} \le r_n\} = n | N_T) = e^{-\mu} \frac{\mu^n}{n!} \quad \text{where } \mu \equiv \frac{V_d}{V_T} N_T. \tag{10}$$

We thus show that the number of axonal arbors covering a random position in space can be estimated from a Poisson distribution, with the mean proportional to the number of remaining axonal arbors (i.e. neurons).

#### Concentration of dopamine inside an isolated area

We considered the effects of diffusion in a region where no dopaminergic axonal arbors and terminals are left. Mathematically this corresponds to a situation where a sphere of radius $r$ is surrounded by a large region with a steady state dopamine level $C_0$. Thus, we here wished to solve the diffusion equation in spherical coordinates

$$\frac{\partial C(r,t)}{\partial t} = D\left(\frac{\partial^2 C(r,t)}{\partial r^2} + \frac{1}{r}\frac{\partial C(r,t)}{\partial r}\right), \tag{11}$$

with boundary conditions

$$C(a,t) = C_0, \quad C(0,t) = 0, \quad C(r,0) = 0 \quad (r < a). \tag{12}$$

Using a proper substitution and Laplace transformation, one can show that the concentration is described by

$$C(r,t) = C_s \cdot \left(1 + \frac{2a}{\pi r} \sum_{n=1}^{\infty} \frac{(-1)^n}{n} \sin\left(\frac{2\pi r}{a}\right) e^{-D\frac{n^2 \pi^2 t}{a}}\right). \tag{13}$$

However, this does not include the decay of dopamine. In the healthy striatum, this quantity is negligible, but in an isolated area, where most arbors are lost, this becomes important. We therefore solved

$$\frac{\partial C(r,t)}{\partial t} = D\left(\frac{\partial^2 C(r,t)}{\partial r^2} + \frac{1}{r}\frac{\partial C(r,t)}{\partial r}\right) - kC(r,t). \tag{14}$$

The solution is obtained from the above result, through the relation

$$C(r,t) = k \int_0^t C_1(r,t') e^{-kt'} dt' + C_1(r,t) e^{-kt}. \tag{15}$$

Hence, the dopamine concentration in an isolated area can be described by

$$C(r,t) = C_s + \frac{2kC_s a}{\pi r}\sum_{n=1}^{\infty}\frac{(-1)^n}{n}\cdot\sin\left(\frac{n\pi r}{a}\right)\left(\frac{k}{X}\left(1-e^{-Xt}\right)+e^{-Xt}\right) \qquad \text{with } X \equiv \frac{Dn^2\pi^2}{a^2}. \tag{16}$$

#### Voronoi tessellation

In $X$ space, with a $d(x, P_i)$ distance function, and $P_i$ points, we defined the most isolated area as the largest sphere one can draw, without meeting one of the N points. We therefore searched for the center of this sphere, termed the mark, which has the largest distance to the nearest point. Mathematically we formulated the mark $x^*$ as:

$$x^* = \max\Big\{\min\{d(x, P_j)\}\Big\}$$
$$x^* = \min\Big(x \in X|\partial_x(\min\{d(x, P_j)\}) = 0\Big).$$

Since we searched for a minimum in 3D, but filled with local minima created by all points, finding this mark is a surprisingly tedious task, and using brute force Monte Carlo techniques takes much computational time and gives low precision. To enhance both calculation time and precision, we employed the concept of Voronoi Diagrams. A Voronoi cell, $R_k$ is defined as the set of all points, whose distance to $P_k$ is not greater than their distance to any of the other sites $P_j$. Thus, the formal definition of a Voronoi diagram is

$$R_K = \Big\{x \in X|d(x, P_k) \leq d(x, P_j) \quad \text{for all } k \neq j\Big\}.$$

Using this definition, the most isolated point must be on the edges where different Voronoi cells intersect, and we thus searched for the most isolated points in the set of all Voronoi cell edges. Using this technique, we found all the isolated points inside the area. However, since some of the edges meet outside the circle denoting striatum space, the most isolated areas on the boundary are not found and we therefore searched for points on the boundary as well. Comparing these distances to the distribution found in Fig. 1h, we could be sure that no regions were more isolated than the ones found by these measures.

#### Defining contiguous arbor classes

We considered the communication classes of the dopaminergic network - termed contiguous arbor classes (CACs) - by defining that two neurons $i$, $j$ communicate if there is a possible path from $i$ to $j$. This means that they do not need to be directly functionally connected, but they should indirectly be able to transmit information between each other. If the dopaminergic neurons communicated we defined the network to be irreducible. To find the CACs we devised the following algorithm:

- pick neuron $n_1$ and put it in set $S_1$;
- find all its axonal arbor neighbors and put these in a transient set C $= \{n_j, \ldots, n_k\}$;
- pick the first element of $C$, put it in $S_1$, remove it from C, and find all its neighboring arbors and put these in C;

- pick the next element of $C$ and repeat the algorithm. When $C$ is empty, set $S_1$ is a collection of all dopaminergic neurons in CAC 1.

- After this, take $n_2$. If $n_2 \in S_1$ go to $n_3$. Otherwise create set $S_2$, put all the connections of $n_2$ in $C$, and repeat the algorithm as above.

#### Removing dopaminergic neurons

To simulate denervation of the striatum we compared three algorithms that emulate three molecular mechanisms. Fundamentally, all the algorithms were event-driven Gillespie algorithms (Gillespie 1977), meaning that the time of the next event was chosen by

$$t_{next} = t - \frac{\ln(\nu)}{\sum_{i=1}^{N} \lambda_i},$$

while the neuron k, that was chosen to die, was selected by fulfilling the criteria

$$\frac{\sum_{i=1}^{k} \lambda_i}{\sum_{j=1}^{N} \lambda_j} \leq \nu' \leq \frac{\sum_{i=1}^{k+1} \lambda_i}{\sum_{j=1}^{N} \lambda_j}.$$

Here $\nu, \nu'$ are uniformly distributed random numbers and the defined rates $\lambda_i$ differed fundamentally between the three models and are explained below.

- Random denervation: At time t=0 all neurons have the same probability to die. This means that they all have a constant rate, and therefore $\lambda_i = \lambda_0^r$.

- Prion-like denervation: At time t=0 one neuron is infected and dies with rate $\lambda_0^p$. This updates the time, and before it dies, it passes on the infection to two neighboring neurons, if two neighbors that are not yet infected exist. Next, one of these will be chosen, so $t_{next} = t - \frac{\ln(\nu)}{\sum_{i=1}^{N_I} \lambda_i}$ with $\nu$ being a uniform random number and $N_I$ being the number of infected neurons.

- Stress-induced denervation: At time t=0 all neurons have a rate to die depending on the number of overlapping axonal arbors $n_i$. Therefore, we define the rate as a logistic function given by $\lambda_i = \lambda_0^s \frac{e^{-\beta(n_i-\gamma)}}{1+e^{-\beta(n_i-\gamma)}}$. Here $\gamma$ is the threshold value, chosen to be 15 since this is approximately half the number in the healthy striatum, and $\beta$ is the steepness, chosen to be 10.

#### Averaged-Neuron SPN model

We consider the membrane potential ($V_m$) given by:

$$dV_m = dt\left(-\frac{1}{AC}\left(\sum I_{Ext}\right) - \frac{1}{C}\left(\sum I_{Int}\right)\right) + D \cdot W_r, \tag{17}$$

with the Intrinsic (Int) and Extrinsic (Ext) conductances that correspond to

$$\text{Int} \in [Leak, Na_V, K_V, K_{A-type}, K_{SI}, Ca_V, K_{Ca}, NaP, K_{IR}]$$
$$\text{Ext} \in [NMDA, AMPA, GABA_A].$$

The unit on the left-hand side of Eq. (17) is mV, whereas the time is measured in ms; on the other hand, the extrinsic and intrinsic currents are respectively in units of µA and nA, which means that the first should be multiplied by $10^3$. However, given that A is measured in $cm^2$, to have consistency with the overall choice of units, $I_{Ext}$ should just be multiplied by 10. The term $D \cdot W_r$ is a stochastic noise term, where $D$ is a parameter quantifying the magnitude of the noise, and $W_r$ is a classical Wiener process.

#### Intrinsic ion channel conductances

In the following we report a list of the intrinsic ion channel conductances considered.

- Leak channel:

$$I_{Leak} = g_{Leak}(V - V_{Leak}) \tag{18}$$
$$g_{Leak} = 0.035\,73\,\mathrm{mS\,cm^{-2}} \quad V_{Leak} = \frac{RT}{zF}\ln\left(\frac{p_K[K]_o + p_{Na}[Na]_o + p_{Cl}[Cl]_i}{p_K[K]_i + p_{Na}[Na]_i + p_{Cl}[Cl]_o}\right)$$
$$R = 8.314\,472\,\mathrm{J\,K^{-1}\,mol^{-1}} \quad T = 310\,\mathrm{K} \quad z = valence_{ion} \quad F = 9.648\,533\,99 \times 10^4\,\mathrm{C\,mol^{-1}}$$

- Voltage-gated sodium channel:

$$I_{Na_V} = g_{Na_V} m_{Na_V}^3 h_{Na_V}(V - V_{Na}) \tag{19}$$
$$m_{Na_V} = \frac{\alpha_m}{\alpha_m + \beta_m} \qquad \begin{cases} \alpha_m = 0.1\frac{V+33}{1-e^{-(V+33)/10}} \\ \beta_m = 4e^{-(V+53.7)/12} \end{cases}$$
$$\dot{h}_{Na_V} = 4(\alpha_h(1 - h_{Na_V}) - \beta_h h_{Na_V}) \qquad \begin{cases} \alpha_h = 0.07e^{-(V+50)/10} \\ \beta_h = \frac{1}{1+e^{-(V+20)/10}} \end{cases}$$
$$g_{Na_V} = 2.4637\,\mathrm{mS\,cm^{-2}} \quad V_{Na} = \frac{RT}{zF}\ln\left(\frac{[Na]_o}{[Na]_i}\right)$$

- Voltage-gated potassium channel:

$$I_{K_V} = g_{K_V} n_K^4 (V - V_K) \tag{20}$$
$$\dot{n}_{K_V} = 4(\alpha_h(1 - h_{K_V}) - \beta_h h_{K_V}) \qquad \begin{cases} \alpha_n = 0.01\frac{V+34}{1-e^{-(V+34)/10}} \\ \beta_n = 0.125e^{-(V+44)/25} \end{cases}$$
$$g_{K_V} = 2.918\,68\,\mathrm{mS\,cm^{-2}} \quad V_K = \frac{RT}{zF}\ln\left(\frac{[K]_o}{[K]_i}\right)$$

- Fast A-type potassium channel:

$$I_{A-type} = g_{A-type} m_{A-type}^3 h_{A-type}(V - V_K) \tag{21}$$
$$m_{A-type} = \frac{1}{1 + e^{-(V+50)/20}}$$
$$\dot{h}_{A-type} = \frac{h_{A-type\infty} - h_A - type}{\tau_{hA-type}}$$
$$h_{A-type\infty} = \frac{1}{1 + e^{(V+80)/6}}$$
$$g_{A-type} = 2.2259\,\mathrm{mS\,cm^{-2}} \quad \tau_{hA-type} = 15\,\mathrm{ms} \quad V_K = \frac{RT}{zF}\ln\left(\frac{[K]_o}{[K]_i}\right)$$

- Slowly inactivating potassium channel:

$$I_{K_{SI}} = g_{K_{SI}} m_{K_{SI}}(V - V_K) \tag{22}$$
$$m_{K_{SI}} = \frac{h_{m_{K_{SI}\infty}} - m_{K_{SI}}}{\tau_{mK_{SI}}}$$
$$m_{K_{SI}\infty} = \frac{1}{1 + e^{-(V+34)/6.6}}$$
$$\tau_{mK_{SI}} = \frac{8}{e^{-(V+55)/30} + e^{(V+55)/30}}$$
$$g_{K_{SI}} = 0.035\,013\,5\,\mathrm{mS\,cm^{-2}} \quad V_K = \frac{RT}{zF}\ln\left(\frac{[K]_o}{[K]_i}\right)$$

- Voltage-gated calcium channel:

$$I_{Ca_V} = g_{Ca_V} m_{Ca_V\infty}^2(V - V_{Ca}) \tag{23}$$
$$m_{Ca_V\infty} = \frac{1}{1 + e^{(V+20)/9}}$$
$$g_{Ca_V} = 0.256\,867\,\mathrm{mS\,cm^{-2}} \quad V_{Ca} = \frac{RT}{zF}\ln\left(\frac{[Ca]_o}{[Ca]_i}\right)$$

- Calcium-dependent potassium channel:

$$I_{K_{Ca}} = g_{K_{Ca}} m_{K_{Ca}\infty}(V - V_K) \tag{24}$$
$$m_{K_{Ca}\infty} = \frac{1}{1 + \frac{K_D}{[Ca]_i}^{3.5}}$$
$$\dot{[Ca]}_i = -\alpha_{Ca}(10 \cdot AI_{Ca} + I_{NMDA}) - \frac{[Ca]_i}{\tau_{Ca}}$$
$$g_{K_{Ca}} = 2.349\,06\,\mathrm{mS\,cm^{-2}} \quad K_D = 30\,\mu\mathrm{M} \quad \tau_{Ca} = 121.403\,\mathrm{ms} \quad V_K = \frac{RT}{zF}\ln\left(\frac{[K]_o}{[K]_i}\right)$$

- Persistent sodium channel:

$$\begin{aligned} I_{NaP} =& g_{NaP} m_{NaP\infty}(V - V_{Na}) \\ m_{NaP\infty} =& \frac{1}{1 + e^{-(V+55.7)/7.7}} \\ g_{NaP} =& 0.071\,798\,4\,\mathrm{mS\,cm^{-2}} \quad V_{Na} = \frac{RT}{zF}\ln\left(\frac{[Na]_o}{[Na]_i}\right) \end{aligned} \tag{25}$$

- Inwardly rectifying potassium channel:

$$\begin{aligned} I_{K_{IR}} =& g_{K_{IR}} h_{K_{IR}\infty}(V - V_K) \\ h_{K_{IR}\infty} =& \frac{1}{1 + e^{(V+75)/4}} \\ g_{K_{IR}} =& 0.016\,645\,4\,\mathrm{mS\,cm^{-2}} \quad V_K = \frac{RT}{zF}\ln\left(\frac{[K]_o}{[K]_i}\right). \end{aligned} \tag{26}$$

We included in the model that the cAMP level should affect the gating channels of the model, especially $K_{SI}$, $K_{Ca}$ and $Ca_V$. In all cases, cAMP would stimulate the activity of PKA, which would in turn decrease the efficiency of all three channels. Therefore we modelled the effect of cAMP as an inhibiting effect on these three gating channels by introducing the following three dependencies:

$$\begin{aligned} g_{K_{SI}} &\mapsto g^0_{K_{SI}} \kappa_{K_{SI}}/(\kappa_{K_{SI}} + cAMP) \\ g_{K_{Ca}} &\mapsto g^0_{K_{Ca}} \kappa_{K_{Ca}}/(\kappa_{K_{Ca}} + cAMP) \\ g_{Ca_V} &\mapsto g^0_{Ca_V} \kappa_{Ca_V}/(\kappa_{Ca_V} + cAMP). \end{aligned}$$

Here we have used $\kappa_{K_{SI}} = 1\,\mu\mathrm{M}$, $\kappa_{K_{Ca}} = 1\,\mu\mathrm{M}$ and $\kappa_{Ca_V} = 0.01\,\mu\mathrm{M}$. In the simulations we have assumed a standard value for the gating channel during tonic release, and then stimulated it by a maximal cAMP level during short phasic periods of 0.4 s.

#### Extrinsic ion channel conductances

First of all, the saturating function was defined as

$$f(V) = \frac{1}{1 + e^{-(V-20)/2}}.$$

Below we report a list of the extrinsic ion channel conductances considered.

- AMPA receptor:

$$\begin{aligned} I_{AMPA} =& g_{AMPA} s_{AMPA}(V - V_{AMPA}) \\ \dot{s_{AMPA}} =& 3.48 f(V) - \frac{s_{AMPA}}{\tau_{AMPA}} \\ g_{AMPA} =& 0.513\,425\,\mu\mathrm{S\,cm^{-2}} \quad V_{AMPA} = \frac{RT}{zF}\ln\left(\frac{p_K[K]_o + p_{Na}[Na]_o}{p_K[K]_i + p_{Na}[Na]_i}\right) \end{aligned} \tag{27}$$

- NMDA receptor:

$$
\begin{aligned}
I_{NMDA} =& \frac{1.1}{1.0 + [Mg]_o/8.0\,\text{mM}} g_{NMDA} s_{NMDA} (V - V_{NMDA}) \\
& s_{N\dot{M}DA} = 0.5 x_{NMDA}(1 - s_{NMDA}) - \frac{s_{NMDA}}{\tau_{sNMDA}} \\
& x_{N\dot{M}DA} = 3.48 f(V) - \frac{x_{NMDA}}{\tau_{xNMDA}} \\
& g_{NMDA} = 0.004\,341\,32\,\mu\text{S}\,\text{cm}^{-2} \quad V_{NMDA} = \frac{RT}{zF} \ln\left(\frac{p_K[K]_o + p_{Na}[Na]_o + p_{Ca}[Ca]_o}{p_K[K]_i + p_{Na}[Na]_i + p_{Ca}[Cl]_i}\right)
\end{aligned}
\tag{28}
$$

- $\text{GABA}_A$ receptor:

$$
\begin{aligned}
I_{GABA_A} =& g_{GABA_A} s_{GABA_A} (V - V_{GABA}) \\
& s_{GA\dot{B}A_A} = f(V) - \frac{s_{GABA_A}}{\tau_{sGABA_A}} \\
& g_{GABA_A} = 0.002\,529\,16\,\mu\text{S}\,\text{cm}^{-2} \quad V_{GABA} = \frac{RT}{zF} \ln\left(\frac{[Cl]_i}{[Cl]_o}\right)
\end{aligned}
\tag{29}
$$

To model the effect of increased levels of synaptic barrages, we increased the magnitude of the stochastic noise term ($D$) included in equation (17) for the membrane potential. In particular:

$$
\begin{aligned}
D &= 0.42\,\text{mV in normal conditions,} \\
D &= 0.98\,\text{mV during increased synaptic barrages.}
\end{aligned}
$$

#### Intra- and extracellular ion concentrations

We used the following intra- and extracellular ion concentrations:

$$
\begin{aligned}
&[Na^+]_o = 140\,\text{mM} && [Na^+]_i = 7\,\text{mM} \\
&[K^+]_o = 3.0\,\text{mM} && [K^+]_i = 140\,\text{mM} \\
&[Ca^{2+}]_o = 1.35\,\text{mM} && [Ca^{2+}]_i = (-\alpha_{Ca^{2+}}(10 \cdot AI_{Ca} + I_{NMDA}) - [Ca^{2+}]i/\tau_{Ca^{2+}})\,\mu\text{M} \\
&[Cl^-]_o = 140\,\text{mM} && [Cl^-]_i = 10\,\text{mM} \\
&[Mg^{2+}]_o = 0.8\,\text{mM}
\end{aligned}
$$

#### Izhikevich SPN model

To model SPN firing activity, we used the model proposed by Izhikevich (Izhikevich, 2003). This is a minimal model, where the different parameters can lead to distinct firing dynamics, such as tonic and burst firing. In this model we describe the neuronal firing through two coupled differential equations:

$$\frac{dv}{dt} = 0.04v^2 + 5v + 140 - u + \mathcal{I}$$
$$\frac{du}{dt} = a(bu - v)$$
$$\text{where} \quad b = b_0 + cAMP_{D_I}$$
$$\text{with the reset condition:}$$
$$\text{if } v > 30 \begin{cases} u = u + d \\ v = c \end{cases}$$

In this model, v and u are dimensionless variables, where v represents the membrane potential of the SPN neuron and u represents a membrane recovery variable. This accounts for the activation of $K^+$ and inactivation of $Na^+$ ionic currents. Furthermore, synaptic currents or injected currents are delivered through the variable $\mathcal{I}$, which acts as stochastic noise in the case of synaptic currents.

#### Postsynaptic compensatory mechanisms

We modelled increased numbers of D2 receptors by decreasing $\lambda_2$, proportionally to the compensation parameter $\delta$,

$$\lambda_2^- \mapsto \lambda_2(1-\delta).$$

We modelled enhanced D1 and D2 receptor sensitivity by modifying the parameters $k_{1,2}$,

$$k_{1,2}^- \mapsto k_{1,2}(1-\delta).$$

Finally, we modelled suppressed cAMP degradation in D1 and D2-SPNs by changing the parameters $\delta_{1,2}$,

$$\delta_{1,2}^- \mapsto \delta_{1,2}(1-\delta).$$

None of the three postsynaptic mechanisms tested effectively restored the original (i.e. the healthy) cAMP signals in both types of neurons. Indeed, since denervation results in an incomplete removal of extracellular DA during pauses, increasing the number of D2 receptors, or their sensitivity, leads to the undesired effect of flattening the cAMP peaks in D2-SPNs even more. On the other hand, enhancing the sensitivity of D1 receptors results in increased cAMP levels during both tonic and phasic dopaminergic neuronal firing, therefore not improving the signal. Finally, modelling a suppressed degradation of cAMP appears to restore the cAMP signal during phasic firing in D1-SPNs, although leading to a slower cAMP clearance afterwards, but the same apparoch is not efficient in D2-SPNs.

### 2 Figures

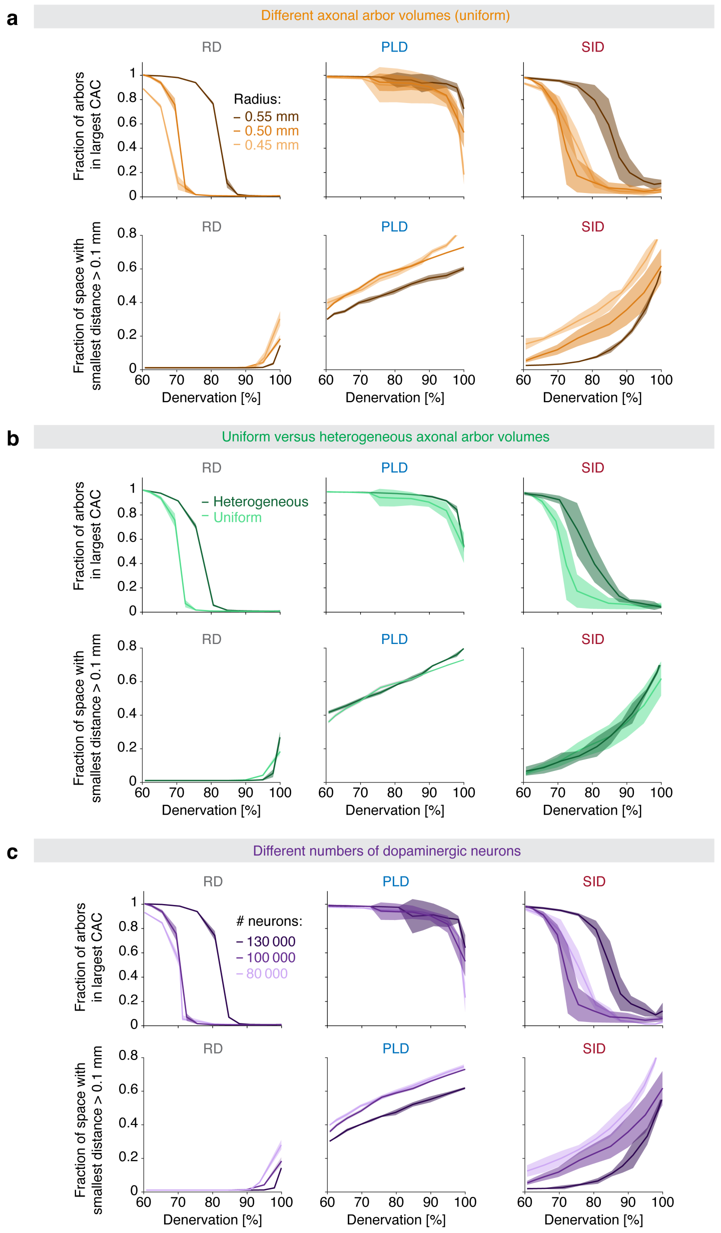


**Supplementary Fig. 1 | Spatial denervation evolutions are robust to changes in key parameters.** **a**, Effects of varying the volume of axonal arbors (radius: 0.45 mm, 0.5 mm or 0.55 mm) uniformly across the arbor population in the three denervation models (random denervation [RD], prion-like denervation [PLD], and stress-induced denervation [SID]). Upper: Fraction of arbors belonging to the largest contiguous arbor class (CAC) as a function of denervation; lower: fraction of striatal space with smallest distance to nearest arbor larger than 0.1 mm (isolated area) as a function of denervation. **b**, Same as in **a**, but with the volume of arbors following a $\delta$ function ($V_n = \delta(V_n - V_0)$) versus heterogeneous distribution so the volume follows a normal distribution ($V_n = V_0 + V_0/10 \times \mathcal{N}(0,1)$). For both cases $V_0 = \frac{4}{3}\pi r_0$ where $r_0$ is the standard radius ($r_0 = 0.5\,\text{mm}$). **c**, Same as in **a**, but for different numbers of dopaminergic neurons in the healthy state (80 000, 100 000 or 130 000 neurons). In **a**–**c**, full line is the mean and shading is the standard deviation.

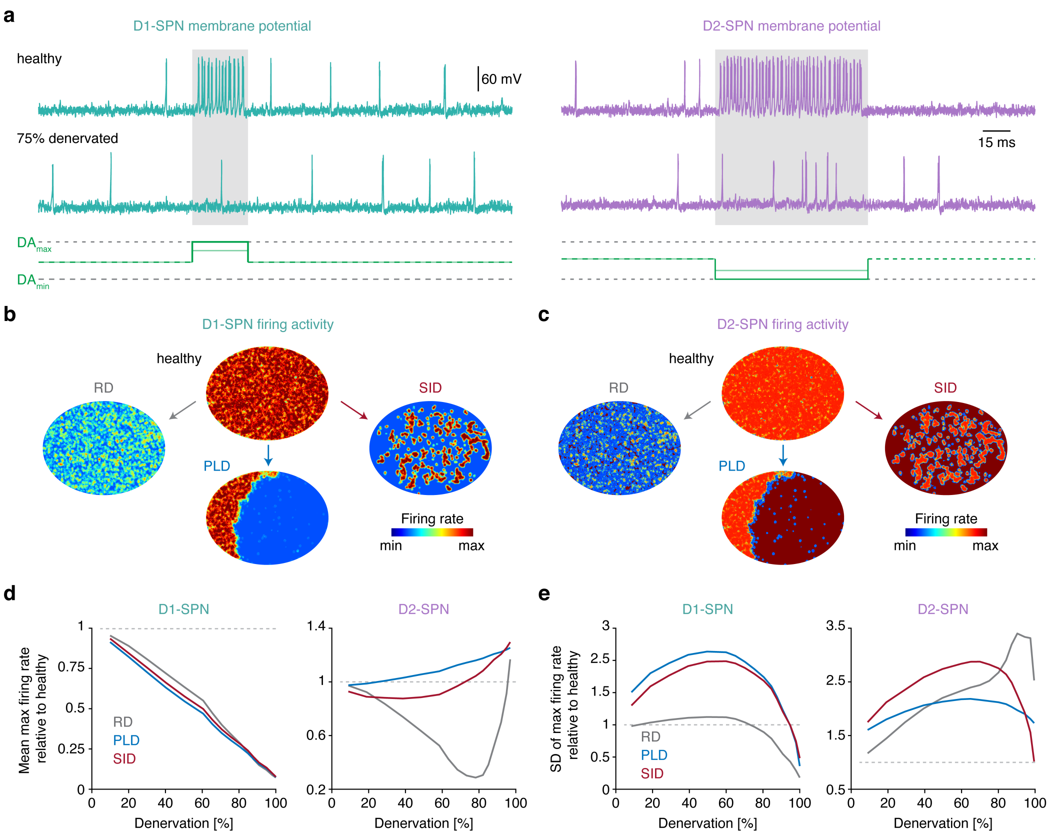


**Supplementary Fig. 2 | Distinct denervation patterns differentially affect local and global striatal projection neuron (SPN) firing activity in the Izhikevich model. a**, Membrane potential of D1-SPN (left) and D2-SPN (right) in the healthy and 75 % denervated striatum in response to dopamine (DA) signaling, modelled using the Izhikevich model. **b–c** Maximal firing activity of D1- and D2-SPNs across space in the healthy and 75 % denervated striatum for the three denervation patterns. **d–e** Spatial mean and standard deviation of maximum firing activity in D1- and D2-SPNs as a function of denervation.

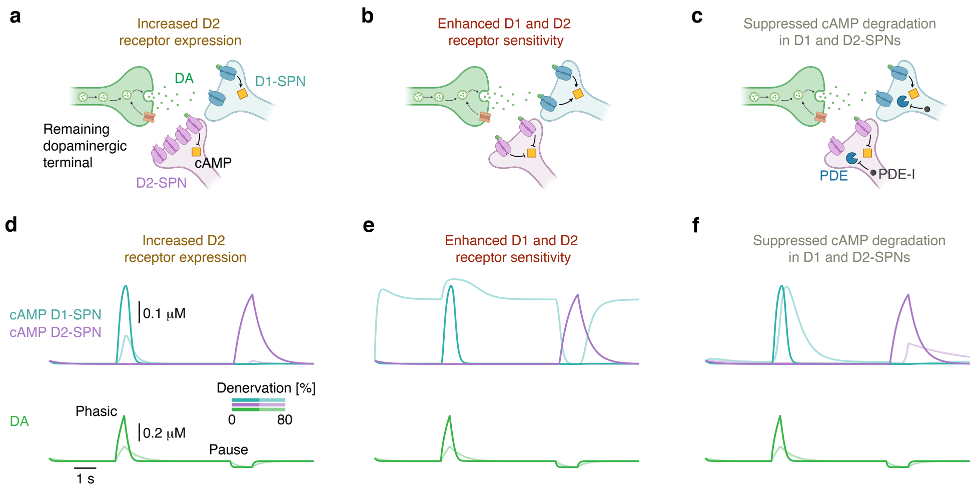


**Supplementary Fig. 3 | Three distinct postsynaptic mechanisms fail to preserve dopamine (DA) signaling in the denervated striatum. a–c** Diagrams of the modelled postsynaptic compensatory mechanisms: increased D2 receptor expression (**a**), enhanced D1 and D2 receptor sensitivity (**b**), and suppressed cyclic adenosine monophosphate (cAMP) degradation in D1 and D2 striatal projection neurons (SPNs) mediated by, for example, a phosphodiesterase (PDE) inhibitor (PDE-I) (**c**). **d–f** Example traces showing cAMP in D1- and D2-SPNs as a function of DA signaling at 0 % and 80 % denervation in the different postsynaptic compensation models.

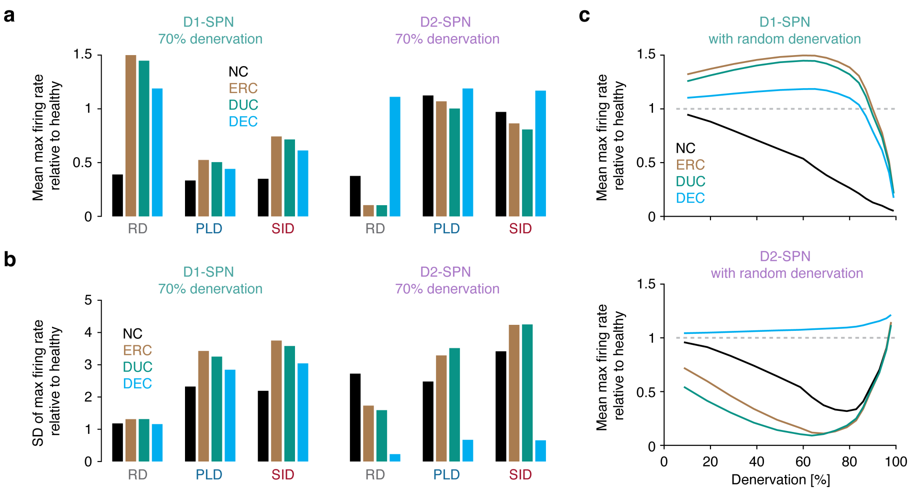


**Supplementary Fig. 4 | A dual presynaptic compensation strategy preserves striatal projection neuron (SPN) firing activity in the Izhikevich model in spite of severe denervation. a–b** Spatial mean and standard deviation of maximum firing activity in D1- and D2-SPNs as a function of denervation pattern (random denervation [RD], prion-like denervation [PLD], and stress-induced denervation [SID]) and compensation model (enhanced release compensation [ERC], decreased uptake compensation (DUC), and dual enhanced compensation [DUC]). **c**, Spatial mean of maximum firing activity in D1- and D2-SPNs as a function of denervation and compensation model in the randomly denervated striatum.

### 3 Tables

#### Supplementary Table 1. Parameters in the model

| | |
|---|---|
| Size of subvolume | $(10 \times 10 \times 10)\,\mu m^3 = 1000\,\mu m^3$ |
| Size of striatum (axes) | a = 0.3 cm |
| | b = 1.5 cm |
| | c = 2.1 cm |
| Size of striatum (volume $V_T$) | $4\pi/3\,(0.3 \times 1.5 \times 2.1)\,cm^3 = 3.96\,cm^3$ |
| Axonal arbor of neuron (radius $r_d$) | 0.5 mm |
| Axonal arbor of neuron (volume $V_d$) | $4\pi/3\,(0.5 \times 0.5 \times 0.5)\,mm^3 = 0.54\,mm^3$ |
| Number of neurons projecting into healthy striatum ($N_T$) | 100 000 |
| Diffusion coefficient in striatum | $380\,\mu m^2\,s^{-1}$ |
| **Dopaminergic neuron firing parameters** | |
| $\Delta$ (release per terminal) | 0.0025 µM |
| $V_M$ (uptake per terminal) | $0.041\,\mu M\,s^{-1}$ |
| $\nu$ (firing frequency tonic) | 4 Hz |
| $\nu$ (firing frequency pause) | 0 Hz |
| $\nu$ (firing frequency phasic) | 15 Hz |
| $\delta$ (degradation of dopamine) | $0.04\,s^{-1}$ |
| $K_M$ (Michaelis-Menten parameter) | 0.21 µM |
| **cAMP production parameters** | |
| $\alpha$ (constant production) | $0.001\,\mu M\,s^{-1}$ (D1-SPN) |
| | $0.001\,\mu M\,s^{-1}$ (D2-SPN) |
| $\lambda$ (receptor stimuli dependent production) | $5.0\,\mu M\,s^{-1}$ (D1-SPN) |
| | $1.0\,\mu M\,s^{-1}$ (D2-SPN) |
| $\kappa$ (affinity) | 0.0025 µM (D1-SPN) |
| | 0.25 µM (D2-SPN) |
| $h$ (Hill coefficient) | 4 (D1-SPN) |
| | 4 (D2-SPN) |
| $\delta$ (active cAMP degradation) | $10\,s^{-1}$ (D1-SPN) |
| | $2\,s^{-1}$ (D2-SPN) |
| **Izhikevich SPN model parameters (dimensionless units)** | |
| a (decay rate for u) | 0.05 |
| b (sensitivity) | 0.2 |
| c (reset value for v) | $-45$ to $-30$ |
| d (reset value for u) | 5 |
